# Functional Identification of Language-Responsive Channels in Individual Participants in MEG Investigations

**DOI:** 10.1101/2023.03.23.533424

**Authors:** Mathias Huybrechts, Rose Bruffaerts, Alvince Pongos, Cory Shain, Benjamin Lipkin, Matthew Siegelman, Vincent Wens, Martin Sjøgård, Idan Blank, Serge Goldman, Xavier De Tiège, Evelina Fedorenko

## Abstract

Making meaningful inferences about the functional architecture of the language system requires the ability to refer to the same neural units across individuals and studies. Traditional brain imaging approaches align and average brains together in a common space. However, lateral frontal and temporal cortices, where the language system resides, is characterized by high structural and functional inter-individual variability, which reduces the sensitivity and functional resolution of group-averaging analyses. This issue is compounded by the fact that language areas lay in close proximity to regions of other large-scale networks with different functional profiles. A solution inspired by visual neuroscience is to identify language areas functionally in each individual brain using a ‘localizer’ task (e.g., a language comprehension task). This approach has proven productive in fMRI, yielding a number of robust and replicable findings about the language system. Here, we extend this approach to MEG. Across two experiments (one in Dutch speakers, n=19; one in English speakers, n=23), we examined neural responses to the processing of sentences and a control condition (nonword sequences). In both the time and frequency domains, we demonstrated that the topography of neural responses to language is spatially stable within individuals but varies across individuals. Consequently, analyses that take this inter-individual variability into account are characterized by greater sensitivity, compared to the group-level analyses. In summary, similar to fMRI, functional identification within individuals yields benefits in MEG, thus opening the door to future investigations of language processing including questions where whole-brain coverage and temporal resolution are both critical.

## 1. Introduction

The functional architecture of the human language network is broadly consistent across individuals (Lipkin et al., 2022). However, the precise topography of this network varies substantially even within homogenous groups of neurotypical adults (Fedorenko et al., 2010). Developing research methods that take these inter-individual differences into account by identifying functional areas in individual participants—an approach known as ‘functional localization’—has proven vital in cognitive neuroscience across domains and has been shown to increase sensitivity, functional resolution, accurate effect size estimation, and interpretability (Saxe et al., 2006; Nieto-Castañón & Fedorenko, 2012; Fedorenko, 2021; see Brett et al., 2002 for an early discussion of these issues). So far, functional localizer paradigms have been mostly used in fMRI research (Kanwisher et al., 1997; Saxe & Kanwisher, 2003; Fedorenko et al., 2010; Baldauf & Desimone, 2014), although a handful of studies have applied a similar approach in other recording modalities, including electrocorticography (ECoG) (Cogan et al., 2014; Fedorenko et al., 2016; Regev et al., 2024) and functional near-infrared spectroscopy (fNIRS) (Powell et al., 2018; Y. Liu et al., 2022; Paranawithana et al., 2024). With respect to magnetoencephalography (MEG), functional localization has predominantly been applied in studies of visual processing (J. Liu et al., 2002; De Vries & Baldauf, 2019), but has also shown potential in the language domain with for instance applications in presurgical mapping and inferring the role of different neural frequency bands during language comprehension (Lam et al., 2016; Prystauka & Lewis, 2019; Papanicolaou, 2023).

Numerous studies over the last decade have provided evidence that the language network can be delineated in a robust and replicable way at the individual-subject level using a contrast between language comprehension and a perceptually matched control condition in fMRI (Fedorenko et al., 2010; Mahowald & Fedorenko, 2016; Braga et al., 2020; Fedorenko et al., 2024; Lee et al., 2024) (see (Lipkin et al., 2022) for data in > 800 individuals). Importantly, the language localizer contrast (language > perceptually matched control condition) has been shown to be robust to input modality (written, spoken, or signed) (Fedorenko et al., 2010; Scott et al., 2017; Richardson et al., 2020; Lee et al., 2024), stimulus content (hand-crafted sentences, sentences extracted from a corpus, or connected passages (Scott et al., 2017)), language (Malik-Moraleda, Ayyash et al., 2022), and the presence or absence of an active task (Fedorenko et al., 2010; Diachek et al., 2020; Ivanova et al., 2020). Furthermore, this network has been shown to be strongly selective for linguistic input (Fedorenko et al., 2011; Monti et al., 2012; Amalric & Dehaene, 2018) (see e.g., (Fedorenko & Blank, 2020) for a review). This broad generalization across paradigm variations and selectivity for language processing jointly suggest that the language brain areas support specifically linguistic computations. Indeed, evidence from dozens of studies has implicated these areas in lexical access, syntactic structure building, and semantic composition during both comprehension and production (e.g. Pallier et al., 2011; Hu, Small et al., 2023; Shain, Kean et al., 2022, 2024; Giglio et al., 2024).

Although fMRI investigations of the language network have yielded important findings, fMRI’s poor temporal resolution limits its use for research questions where timing information is critical. Intracranial recordings provide an incredible opportunity to obtain high-spatial and high-temporal resolution data with high signal-to-noise ratio, but the approach is inherently limited with respect to both the population and the sparse brain coverage. In contrast, magnetoencephalography (MEG) enables whole-brain non-invasive measurements of neural activity in typical brains at a millisecond-level resolution. Here, using data from two independent datasets (across two languages: English and Dutch), we establish the feasibility of identifying language-responsive sensors at the individual level in MEG investigations, and provide evidence that this approach is more sensitive than the standard brain-averaging approach.

## 2. Methods

### 2.1 Participants

We recruited 42 healthy young volunteers: 19 native Dutch speakers (18 female, between 19 and 29 years old, mean 23.4 years old) and 23 native English speakers (between 19 and 53 years old, mean 26.7 years old). The study was approved by the local Ethics Committees. All participants provided written informed consent in accordance with the Declaration of Helsinki.

### 2.2 Experimental Design

In identifying the language network, we use a design that has been extensively validated in fMRI (first used in (Fedorenko et al., 2010)), namely, a sentence reading task. Using fMRI, the sentences>nonwords sequences contrast has been shown to robustly and reliably identify the language-selective network (Lipkin et al., 2022; Malik-Moraleda, Ayyash et al., 2022). Participants performed the experiment in their native language. Materials can be downloaded from https://osf.io/vc2bw/. For the sentence condition, 80 12-word-long sentences were constructed in English using a variety of syntactic structures and covering a wide range of topics. The sentences were translated into Dutch, with minimal changes to obtain 12-word-long sentences. For the nonwords condition, care was taken to minimize low-level differences in the phonological make-up of the stimuli compared to the sentence condition. In English, the content words (noun, verb, adjective, adverb) of the sentence condition were syllabified to create a set of syllables that could be re-combined in new ways to create pronounceable nonwords. For syllables that formed real words of English, a single phoneme was replaced (respecting the phonotactic constraints of English) to turn the syllable into a nonword. The syllables were then recombined to create nonwords matched for length (in syllables) with sentence condition. In Dutch, nonwords were derived from the content words in the sentence condition by means of the pseudoword generator Wuggy (https://github.com/crr-ugent/wuggy). This program generates nonwords that match the original word in subsyllabic structure and respects the transition frequencies specific to Dutch (Keuleers & Brysbaert, 2010).

To help participants stay engaged, we used a memory probe version of the language localizer task in the Dutch version, where participants are asked to decide (by pressing one of two buttons) whether a word or nonword, presented at the end of each sentence or nonword list, was in the preceding trial. Probes were restricted to content words in the sentence condition and nonwords in the nonwords condition. Half of the trials required a positive response. In the English version, a button-press icon was presented at the end of each sentence or nonword list where participants are asked to press a button. This difference between the Dutch and English version was introduced to reduce the duration of the English experiment by several minutes, in order to combine the paradigm with other, unrelated studies. As noted above, previous fMRI work demonstrated that the sentences>nonwords localizer contrast is robust to different tasks including passive viewing (Fedorenko et al., 2010; Diachek et al., 2020).

The words/nonwords were presented one at a time in a rapid serial visual presentation paradigm at a fixed rate per word/nonword (385 ms per (non)word in Dutch, 400 ms in English). Each word/nonword was presented in the center of the screen in capital letters without punctuation and appeared immediately after the previous (non)word without a blank transient screen. Then, at the end of the trial, the memory probe or button-press icon was presented, followed by a variable interval, between 0.5s and 2.25s. Participants could respond any time after the probe or button-press icon appeared and until the next trial started. The experiment lasted ∼20 minutes. Five participants in the English dataset were scanned using a similar paradigm but with a different set of materials (8-word-long sentences and 8-nonword-long sequences). Previous fMRI work has shown that the sentences>nonwords localizer contrast is robust to such variation (Fedorenko et al., 2010).

### 2.3 MEG Data Acquisition

Continuous MEG data were recorded using a whole-head 306 channel (102 magnetometers, 204 planar gradiometers) TRIUX system (MEGIN, Espoo, Finland) either at the HUB - Hôpital Erasme (Brussels, Belgium) or at the Martinos Imaging Center (McGovern Institute for Brain Research at MIT, Cambridge, MA, USA). Participants were tested in an upright seated position (68° recline). Four head-position indicator (HPI) coils were used to record the head position within the MEG helmet every 200 ms. The participant’s head shape was digitally recorded by means of a 3D digitizer (Fastrak Polhemus, Inc., Colchester, VA, USA) along with the position of the HPI coils and fiducial points (nasion, left and right periauricular). MEG signals were recorded at a sampling rate of 1000 Hz with on-line band filter between 0.1 and 330 Hz.

### 2.4 MEG Data Processing

Initial preprocessing of the raw data used MaxFilter version 2.2 (MEGIN, Espoo, Finland): temporal signal space separation (Taulu et al., 2003) was applied to remove noise from external sources and from HPI coils for continuous head-motion correction (correlation threshold: 0.98, 10 s sliding window), and to virtually transform data to a standardized head position. The latter facilitates comparison across experiments. MaxFilter was used to automatically detect and virtually reconstruct noisy channels. Further preprocessing was performed using MNE-Python version 1.8.0 (Gramfort, 2013). Preprocessing consisted of high-pass filtering at 0.1 Hz, low-pass filtering at 300 Hz and notch filtering at 50Hz or 60Hz (with respective higher harmonics) depending on the line frequency. Data were downsampled to 500 Hz and epoched at the level of the sentence or nonword list (500ms before onset of the first (non)word until 400 ms after onset of the last (non)word). Visual inspection of all epochs was performed, and epochs with clear artefacts were marked as bad and excluded from further analysis. Denoising was performed by visually identifying and removing ICA components consistent with eye blinks and cardiac artefacts from a total of 60 components extracted using fastICA (Hyvarinen, 1999) with default parameters on the concatenated epochs (excluding bad epochs).

A quantitative marker of noise was derived per participant by first averaging the baseline intervals across all sensors for each trial and then calculating the standard deviation of these trial-averaged values across all trials. Here and later in the study, the baseline is defined as the neutral interval of 500 ms prior to trial onset when only a fixation cross is presented. Participants with markers of noise greater than two standard deviations from the average noise marker across participants scanned in the same MEG device were removed from further analysis. This procedure led to the removal of one participant from the English dataset. Two additional participants were removed from the English dataset: one due to excessive motion and one due to the presence of high-amplitude “mu rhythm” apparent upon visual inspection.

Language processing related to the presentation of the word or nonword stimuli is reflected in both the time and frequency domain of the recorded MEG signal (Beres, 2017; Prystauka & Lewis, 2019; Coolen et al., 2024). Analyses that focus on the time domain and rely on event-related potentials have reliably shown effects related to prediction and integration of lexical information during language comprehension (Sun & Luo, 2024). However, such analyses only capture a subset of changes in the signal associated with the presentation of the stimuli. Specifically, oscillatory activity that is not strictly time-or phase-locked to the stimulus will be averaged out in this analytic approach (Maguire & Abel, 2013; Morales & Bowers, 2022). To this end, we analyzed the epochs in the time and frequency domains separately. The spectral analysis offers an alternative, complementary perspective to the time domain analysis for understanding the neural computations associated with language comprehension (Lam et al., 2016).

In the time domain analysis, we grouped each pair of planar gradiometers into a single effective gradiometer derived as their Euclidean norm (hence measuring the amplitude of the field’s tangential gradient, independently of orientation) (Chetail et al., 2018). To facilitate between-participant comparisons, we then calculated the percent signal change at every timepoint in the sentence and nonword conditions for every trial compared to the mean amplitude of its baseline. We omitted the first 4 (non)words in each trial (n=80 per condition) from the analysis in order to focus on the part of the trial where the between-condition differences might be most pronounced (given what is known about the processes related to sentence meaning construction (Pallier et al., 2011; Fedorenko et al., 2016)), although this choice proved not to impact the results in the end (see discussion). Finally, we averaged the percent signal change values across all included word/nonword positions (i.e., the 5th (non)word and all subsequent ones) within a condition to derive a single “amplitude-based” value per channel per participant.

In the spectral analysis, we adopted a similar approach: we compared the bandlimited power of the sentence and nonword trials in the 204 planar gradiometers to that of the baseline (as percent power change) for each neural frequency band. We used five bands: θ (3-8Hz), α (8-13Hz), β (13-30Hz), γ_*low*_(30-60Hz) and γ_*high*_(60-90Hz). We divided the γ-band into a lower and higher frequency component given that previous studies have demonstrated distinctions between these sub-bands or linked effects specifically to one of them (Towle et al., 2008; Fedorenko et al., 2016; Lam et al., 2016; Hashimoto et al., 2017). First, we computed the frequency estimates of the sentence trial epochs and nonword trial epochs (from the 5th (non)word onwards, as in the time domain analyses) and their respective baselines using multitapers with 7 Discrete Prolate Spheroidal Sequences (DPSS). Then, we calculated the percent signal change in the power estimates of the sentence and nonword conditions for every trial compared to the baseline interval for that trial. Finally, we averaged all percent signal change values within each condition, which yielded a single “power-based” percent change value for each channel per participant per frequency band.

### 2.5 Statistical analysis

To test whether MEG allows for a robust identification of language-responsive sensors at the individual participant level, we evaluated the stability of the language responses within individuals across time (i.e., between the odd-vs. even-numbered trials). To do so, we performed three analyses. First, we examined the Spearman correlation in the size of the sentence effect (percent signal change for the sentence condition relative to the baseline) over all channels within each participant across odd- and even-numbered trials and compared this to the correlations between different participants (Wilcoxon rank sum test). Spearman instead of Pearson correlation was used because it is more robust to potential outliers (Rousselet & Pernet, 2012). This analysis asks: if a sensor shows a strong response to sentences in one half of the data, does it also show a strong response to sentences in the other half of the data in the same participant compared to another participant?

Second, we used the data from the odd-numbered trials to define sensors of interest (SOIs) in each participant. SOIs were defined as the 10% of sensors with the highest increase in percentage signal change in the sentence condition relative to the baseline. We then examined the effect size for the sentence condition in the even-numbered trials in these SOIs relative to the baseline and to the effect size for the nonword condition using a signed rank test.

Third, to test whether identification of language-responsive sensors at the individual level is superior to identifying language-responsive sensors at the group level, we compared the results of the second analysis above to a version of the analysis where we performed the same calculation using group-averaged maps (where the responses are averaged across participants from either the English or Dutch dataset in each sensor). In particular, we used the data from the odd-numbered trials to create a group map and defined SOIs as the 10% of sensors with the highest increases in percent signal change in the sentence condition relative to the baseline. We then examined the effect size for the sentence condition in the even-numbered trials in these SOIs for each participant. Critically, unlike in the individual-level analyses, the SOIs in the group analysis are the same for all participants. To compare the effect sizes in SOIs defined individually vs. based on a group-level map, we used a signed rank test.

In all analyses, we employed an exploratory-confirmatory approach to control for false positives. The English dataset served as the exploratory sample, where we identified key neural effects of interest. Then, we tested whether these effects could be replicated in the Dutch dataset using the same statistical analysis. Only results that met predefined confirmation criteria – statistical significance (p<0.05), consistent directionality, and comparable effect size – were considered robust. This approach reduces the likelihood of false positives while enhancing the generalizability of our approach.

## 3. Results

### 3.1 The topography of neural responses to language is stable within but varies between individuals

In the English dataset, the average correlation of the amplitude-based sentence effect sizes across all channels within each participant across odd- and even-numbered trials was 0.61 (s.d. 0.22). The mean correlation between pairs of participants was significantly lower compared to the within-participant correlation (mean rho: 0.16, s.d. 0.24; P < 0.0001; Figure 1A). This finding was confirmed in the Dutch dataset as the average within-participant correlation (0.63 s.d. 0.23) was also significantly higher than the average between-participant correlation (0.25 s.d. 0.26; P<0.0001; Figure 1B). Furthermore, the effect was comparable in size and directionality to that of the English dataset, aligning with the predefined confirmation criteria of the exploratory-confirmatory approach.

**Figure 1:**
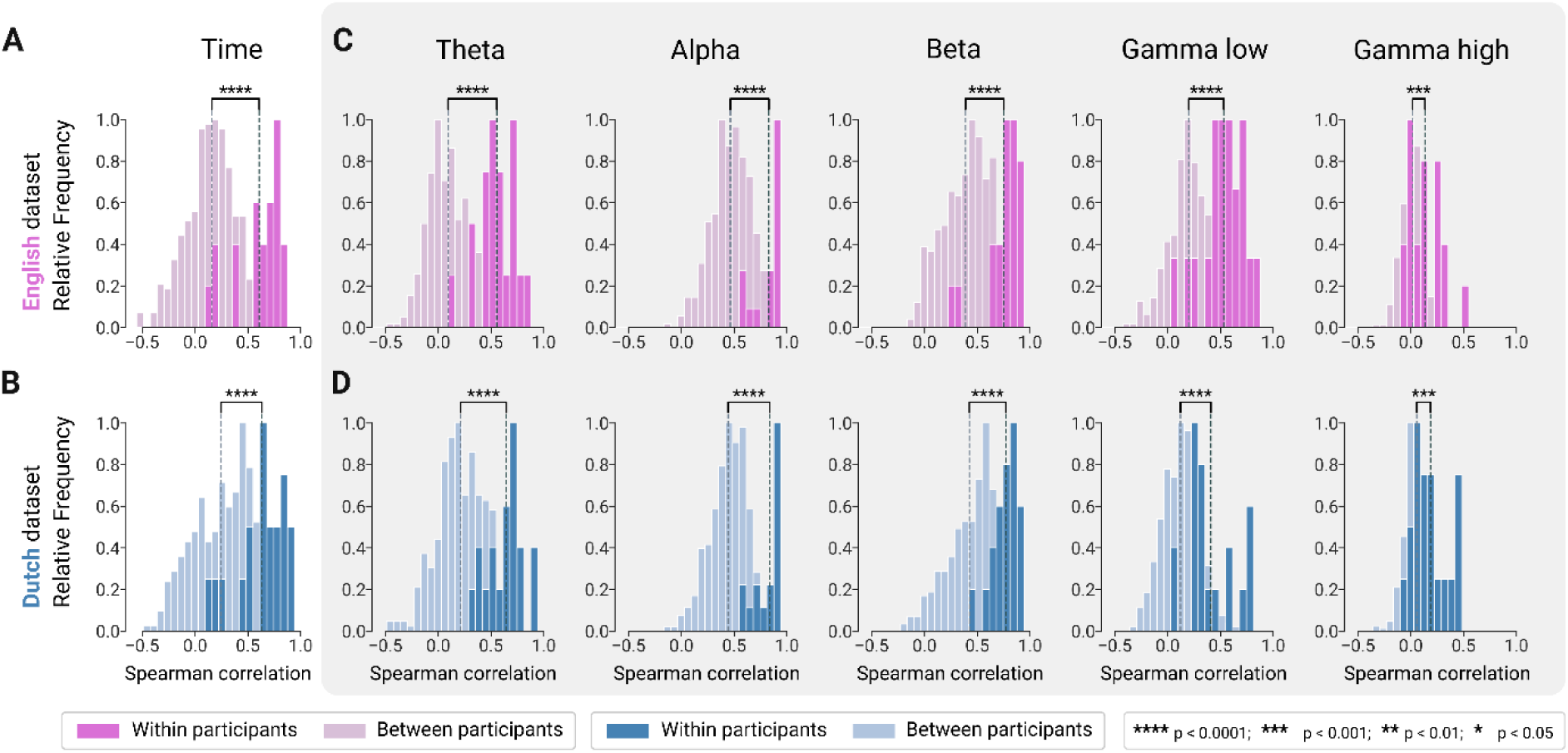
Distribution of the correlation (Spearman) values for the size of the sentence>baseline effect across channels within individuals across odd- and even-numbered trials (darker shades) and between data halves taken from different individuals (lighter shades) in the A) English and B) Dutch datasets in the time domain analysis. Similar distributions for the spectral analysis are represented for the C) English and D) Dutch datasets for each of the neural frequency bands. Relative frequency counts were plotted because of the different number of possible combinations between vs within individuals. Vertical striped lines indicate the mean of each distribution.

In the frequency domain analysis, the average within-participant correlations of the power-based percent changes were significantly higher than the average between-participant correlations (P<0.0001 for all frequency bands apart from high gamma where P<0.001;Figure 1C,D). For each frequency band, similar effect sizes where obtained in the English and Dutch datasets (Table 1).

**Table 1:** Within and between participant correlation values in the English and Dutch datasets for the time domain and each of the neural frequency bands.

| **English** | Time | Theta | Alpha | Beta | Gamma low | Gamma high |
| --- | --- | --- | --- | --- | --- | --- |
| Within | 0.61 | 0.55 | 0.83 | 0.75 | 0.53 | 0.13 |
| correlations | (s.d. 0.22) | (s.d. 0.17) | (s.d. 0.12) | (s.d. 0.17) | (s.d. 0.20) | (s.d. 0.14) |
| Between | 0.16 | 0.09 | 0.46 | 0.39 | 0.20 | 0.02 |
| correlations | (s.d. 0.24) | (s.d. 0.19) | (s.d. 0.18) | (s.d. 0.21) | (s.d. 0.19) | (s.d. 0.10) |
| P value | P < 0.0001 | P < 0.0001 | P < 0.0001 | P < 0.0001 | P < 0.0001 | P < 0.001 |
| **Dutch** | Time | Theta | Alpha | Beta | Gamma low | Gamma high |
| Within | 0.63 | 0.64 | 0.84 | 0.77 | 0.41 | 0.19 |
| correlations | (s.d. 0.23) | (s.d. 0.16) | (s.d. 0.12) | (s.d. 0.12) | (s.d. 0.22) | (s.d. 0.16) |
| Between | 0.25 | 0.21 | 0.44 | 0.42 | 0.12 | 0.06 |
| correlations | (s.d. 0.26) | (s.d. 0.22) | (s.d. 0.17) | (s.d. 0.21) | (s.d. 0.17) | (s.d. 0.11) |
| P value | P < 0.0001 | P < 0.0001 | P < 0.0001 | P < 0.0001 | P < 0.0001 | P < 0.001 |

Inspection confirmed the similarity between the topographies of the signal changes during the sentence condition in the odd- and even-numbered trials at the individual level (Figure 2). When assessing the topography of the language-responsive SOIs that were selected in different participants, inter-individual differences were notable: although some channels were consistently selected in a substantial fraction of participants, other channels were only selected in a small fraction of individuals. In the English dataset, certain sensors were consistently selected in up to 45%, or 9 of the 20 participants in the time domain and up to 75% or 15 of 20 participants in the alpha band. Similarly, in the Dutch dataset, some sensors were consistently selected in up to 42%, or 8 of the 19 participants in the time domain and up to 68% or 13 of 19 participants in the alpha band.

**Figure 2:**
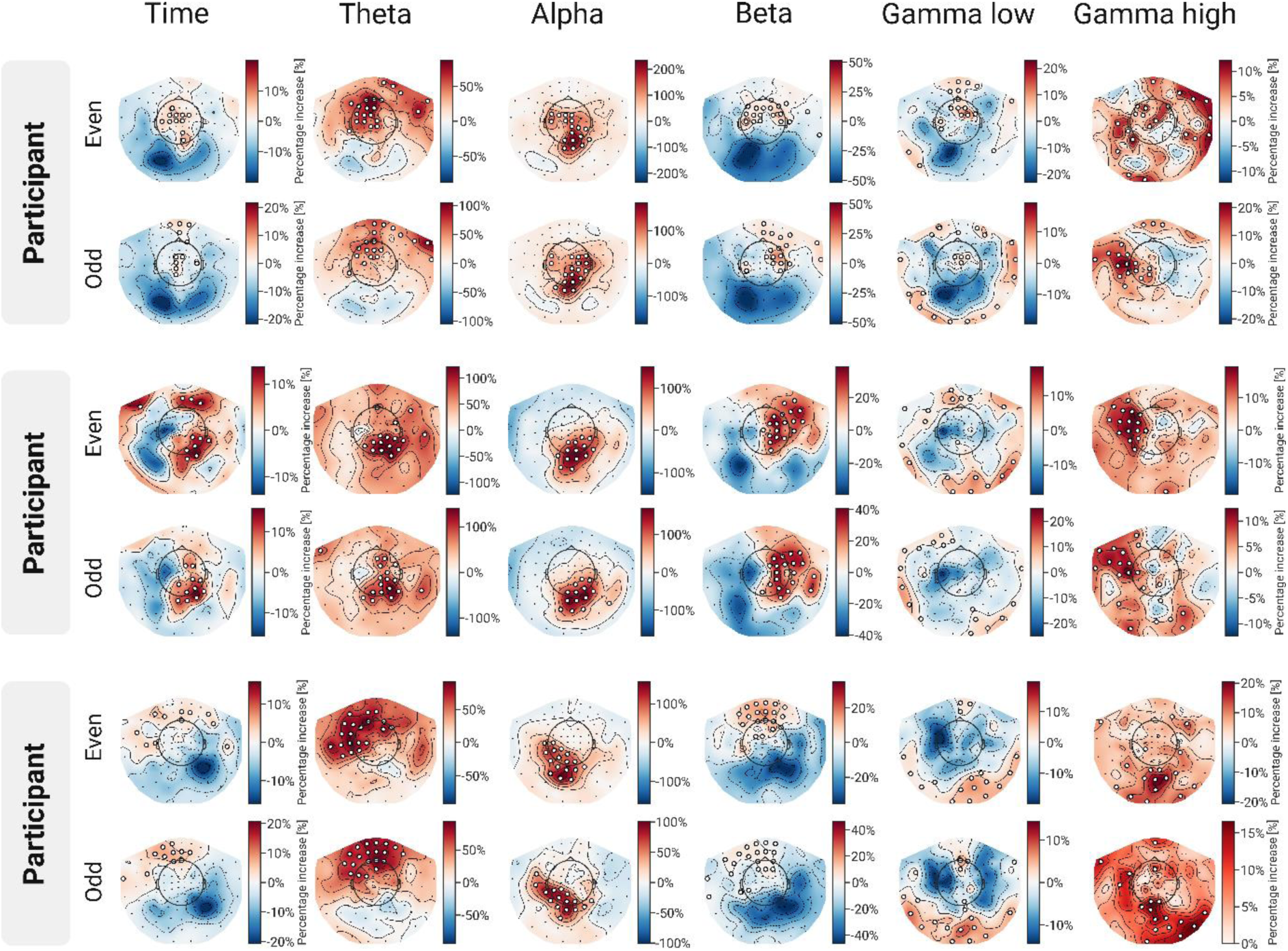
Topographies of the effect sizes (percentage change) during the sentence condition in the odd- and even-numbered trials in time domain and frequency domain in sample participants of the English (top two participants) and Dutch dataset (bottom participant) with selected SOIs indicated by white dots.

### 3.2 Analyses based on individual-level sensors of interest yield more robust responses compared to the group-level analysis

The amplitude-based effect size for the sentence condition in the even-numbered trials with individually defined SOIs (based on the odd-numbered trials) in the English dataset was 3.17% (s.d. 4.04%) signal increase compared to baseline (Figure 3A) and was reliably greater than the effect size for the nonwords condition (P = 0.024). At the level of individual participants, 16 of the 20 showed a sentence > nonwords effect. This was further confirmed in the Dutch dataset where the effect size for the sentence condition in the individually defined SOIs was 5.54% (s.d. 6.04%) signal increase compared to baseline in the even-numbered trials (Figure 3B) and was reliably greater than the effect size for the nonwords condition (P=0.026). At the level of individual participants, 14 of the 19 showed a sentence > nonwords effect (Table 2).

**Figure 3:**
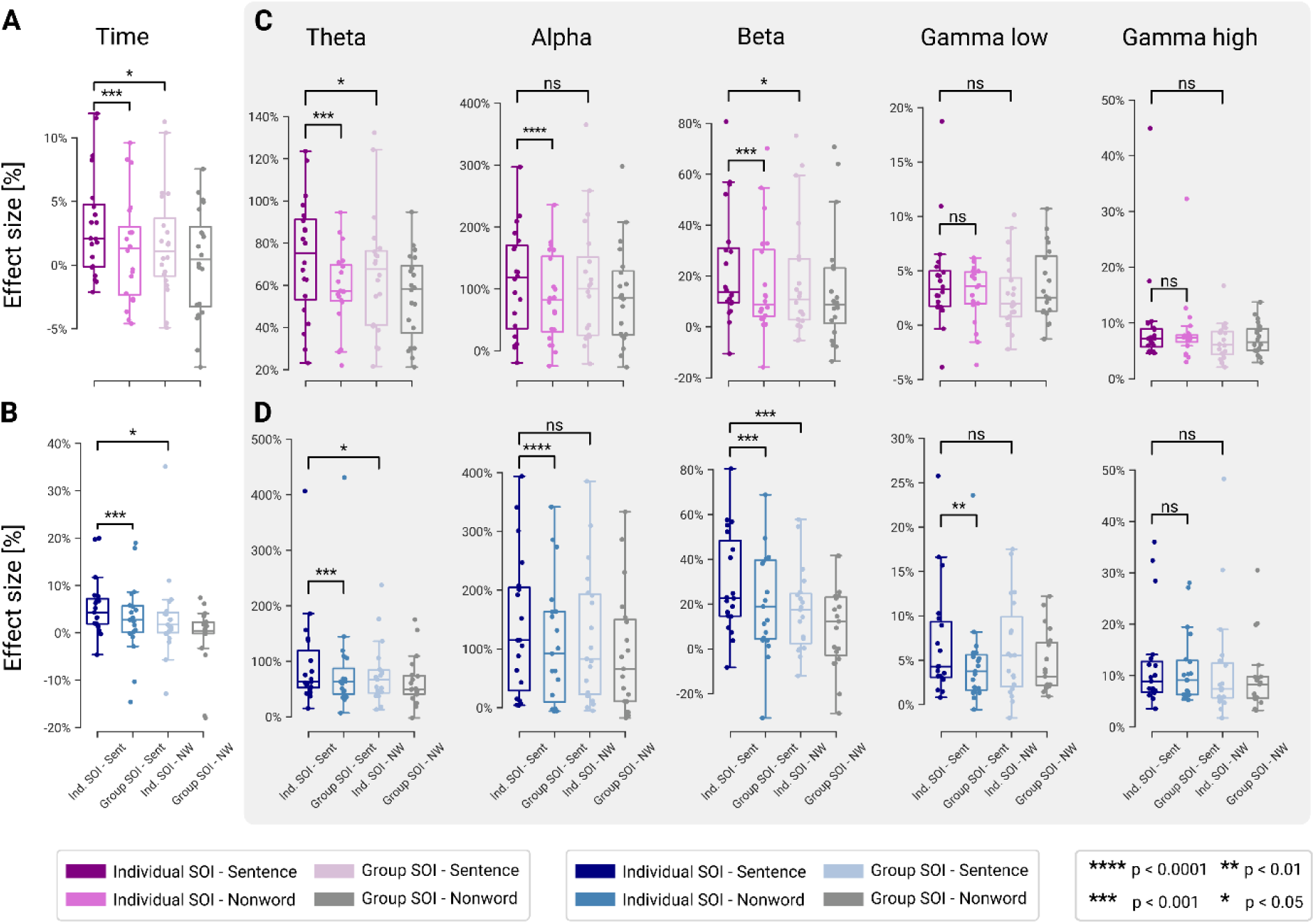
Mean effect sizes (percent signal change) for the sentence and nonwords conditions when the language-responsive sensors of interest (SOIs) are defined at the individual level vs. at the group level in the A) English and B) Dutch datasets for the time domain analysis. Similar results depicted for each of the neural frequency bands in the frequency domain analysis in the C) English and D) Dutch dataset. In all cases, the SOIs are defined using one half of the data (odd-numbered trials) and the response magnitudes are examined in the other half of the data (even-numbered trials).

**Table 2:** (Left) Proportion of participants showing an increase in effect size for the sentence condition over the nonword condition with individually defined SOI (Sentence > Nonword). (Right) The proportion of participants showing an increase in effect size in the sentence condition when SOIs are defined at the individual level versus when SOIs are compared at the group level (Individual SOI > Group SOI). Values indicate the number of significant participants (n) out of the total, with corresponding p-values for English and Dutch datasets.

|  | Individual SOI: Sentence > Nonword |  |  |  | Individual SOI > Group SOI |  |  |  |
| --- | --- | --- | --- | --- | --- | --- | --- | --- |
|  | English |  | Dutch |  | English |  | Dutch |  |
|  | n | p-val | n | p-val | n | p-val | n | p-val |
| Time | 16/20 | 0.024 | 14/19 | 0.026 | 18/20 | < 0.001 | 16/19 | < 0.001 |
| Theta | 16/20 | 0.015 | 16/19 | 0.012 | 16/20 | < 0.001 | 17/19 | < 0.001 |
| Alpha | 12/20 | 0.674 | 14/19 | 0.087 | 19/20 | < 0.0001 | 19/19 | < 0.0001 |
| Beta | 14/20 | 0.014 | 16/19 | < 0.001 | 17/20 | < 0.001 | 15/19 | < 0.001 |
| Gamma Low | 11/20 | 0.294 | 12/19 | 0.441 | 9/20 | 0.596 | 13/19 | 0.006 |
| Gamma High | 14/20 | 0.105 | 11/19 | 0.490 | 10/20 | 0.498 | 12/19 | 0.441 |

In the frequency domain, in the theta and beta bands, power-based signal increases of 72.95% and 23.12% respectively were observed with the individually defined SOIs in the English dataset (Figure 3C). Both were reliably greater than their nonword conditions (P=0.015 for theta and P=0.014 for beta). At the level of individual participants, 16 out of 20 exhibited a greater theta power response to sentences compared to nonword lists, and 14 out of 20 showed an increase in beta power for the same contrast. These findings were replicated in the Dutch dataset where for the theta and beta bands, the increases of the sentence condition over baseline were 96.66% and 29.26%. respectively. Both were reliably greater than the nonword condition with P=0.012 for the theta band and P<0.001 for the beta band. Other frequency bands also showed increased levels of power for the sentence condition compared to baseline but did not demonstrate a reliably greater effect for sentences than for nonwords (Table 2).

To test whether identification of SOIs at the individual level is superior to analysis at the group level, we defined a separate set of SOIs based on the group-level map for the odd-numbered trials. With the group-level SOIs, we found that the amplitude-based effect size for the sentence condition in the English dataset was significantly smaller compared to the analyses that take inter-individual differences into account (P < 0.001, Figure 3A). At the level of individual participants, 18 of the 20 participants showed a larger amplitude-based effect size in the individually defined SOIs compared to the group-level SOI. This was confirmed in the Dutch dataset where the same result was observed (P<0.001, Figure 3B). At the level of individual participants, 16 of the 19 showed a larger effect size when SOIs are selected at the individual level effect. Similarly, the power in the theta, alpha, and beta bands was significantly lower when inter-individual differences were not taken into account for both the English and Dutch dataset (P<0.001, P<0.0001, P<0.001, Figure 3C,D; Table 2).

## 4. Discussion

In two independent datasets, MEG recordings allowed the identification of language-responsive sensors at the individual-participant level, both in the time and frequency domains. During sentence reading, neural signal changes could be detected within native individual English speakers and reproduced in native Dutch speakers. We showed that in both datasets, the response to this language comprehension task is spatially consistent (over time) within individuals: similar sensors show strong responses during language processing in two halves of the data. Importantly, the response to language is spatially variable across individuals, presumably due to differences in the functional neuroanatomy. The consequence is that analyses that take inter-individual variability into account have greater sensitivity than the traditional group-level analysis. Overall, the results suggest that the functional identification approach, where sensors of interest are defined in individual participants, can yield advantages in MEG, including greater sensitivity, functional resolution, and interpretability.

### 4.1 Individual-level identification of the language network

In line with prior results using the same language localizer task with fMRI (Lipkin et al., 2022) and intracranial recordings (Fedorenko et al., 2016), we observed that analyses that take into account inter-individual variability in the functional neuroanatomy of language-responsive cortex yield higher sensitivity: language responses are reliably higher when the channels are selected at the individual level, compared to the group level. When using a typical fMRI preprocessing and analysis pipeline, voxel-wise neural responses from each participant are warped from the subject space to a common space based on a brain template and functional correspondence is assumed in each voxel. This assumption has long been shown to be flawed, especially when examining cognitive functions supported by the association cortex (Nieto-Castañón & Fedorenko, 2012; Frost & Goebel, 2012; Fedorenko & Blank, 2020). Analysis of resting-state MEG data demonstrated that functional connectivity patterns enable differentiation between different individuals (Da Silva Castanheira et al., 2021) and we here extend the individual-level neural characterization to the language network.

In brain recording approaches, like fNIRS and MEG, where a fixed number of sensors are used, a similar assumption is typically made when data are pooled across participants: that the same sensor is functionally equivalent across individuals. However, because of the inter-individual neuroanatomical differences in cortical thickness, gyrification, and total brain volume, and differences in the placement of the recording channels relative to the brain, the same sensor may capture different underlying sources of neural signals depending on the individual’s anatomy. This variability in the sources captured by the same channels across participants introduces noise when signals are averaged across participants and complicates interpretation. potentially leading to incorrect conclusions (e.g., see (Powell et al., 2018) for a discussion of this issue in fNIRS).

Source modelling allows to map neural signals from the sensors to an individual’s structural brain scan for both fNIRS and MEG. However, group-level analysis after source modelling is associated with the same limitations as group-level analysis performed in a common space using fMRI: specifically, warping the neural signals across participants to a common space wrongly assumes functional equivalence of the same voxel or vertex across participants (Nieto-Castañón & Fedorenko, 2012; Frost & Goebel, 2012; Fedorenko & Blank, 2020). Functional identification in individual participants provides a powerful alternative solution. This approach allows us to circumvent potential differences in the underlying anatomy and focus on the functional responses. Functional identification at the individual level can be performed in sensor space, as we have done here, or in source space. Logistically, a MEG source modelling approach requires additional data acquisition using MRI. Future research will investigate the use of source modelling combined with our functional localizer approach to localize the language-responsive vertices at the individual level.

We established the feasibility of individual-participant functional identification in MEG using an extensively validated language localizer paradigm. Previous work established the replicability of neural responses evoked by language processing using MEG within the same group of individuals (Roos & Piai, 2020). We demonstrated the topographic stability of the language-responsive channels within individuals over time—the critical foundation of individual-level functional localization. The functional identification approach yields greater sensitivity and functional resolution. In particular, with respect to sensitivity: by grouping the selected SOIs, one statistical test can be performed across the ensemble of sensors, avoiding the need for multiple statistical comparisons (Nieto-Castañón & Fedorenko, 2012). Further, effect sizes are more accurately estimated. This greater sensitivity can be helpful in examining subtle effects in some new, critical conditions and has the potential for application in patients with neurological disease, where neural responses may be overall weaker.

These advantages in sensitivity and functional resolution are afforded by the application of the individual-subject analyses in any study regardless of whether a functional localizer task is included. However, if a validated localizer paradigm is used, the study will additionally benefit from greater interpretability. By using the same paradigm to identify the relevant functional subset of the brain across individuals, studies, imaging modalities, and species (in cases of shared cognitive capacities e.g., face processing), we can a) make stronger inferences about the origins of an effect (e.g., the ability to interpret some critical effect as arising within the language-responsive regions), which cannot be done based on anatomy alone because of the inter-individual functional differences, and b) be generally more confident that we are referring to the same system which is critical for knowledge accumulation and for comparing findings across studies and labs. In this way, the functional localization approach aligns with the field’s current focus on robust and replicable science (Poldrack et al., 2017). In addition to affording the ability to refer to the same system across studies, studies that rely on functional localization include an internal replication component because half of the data is used to identify the voxels/sensors of interest and the other half is used to quantify the response magnitudes, similar to what is commonly done in other modalities (Peelen & Downing, 2005; Nieto-Castañón & Fedorenko, 2012; Powell et al., 2018). Thus, the use of a functional identification task yields a replication of the effect in every new participant and study, thus reducing type I errors.

We have so far focused on univariate analyses. However, the combined use of functional identification and multivariate analyses, like multivariate pattern analysis (MVPA), could powerfully enable investigations of fine-grained meaning and structure representations within the language network. For example, with respect to semantic knowledge, explicit quantitative models reflecting single-word and contextualized semantic knowledge can be used to test hypotheses about how the brain processes meanings extracted from linguistic input (Bruffaerts et al., 2019). Focusing on language-responsive sensors can increase both the sensitivity of such an analysis and help interpret the observed effects.

### 4.2 Using MEG to probe linguistic computations

We here advocate individual-level functional identification as an approach to achieve greater sensitivity and interpretability when investigating the language network using MEG. The present work introduces this approach using neural signals in sensor space. When dealing with well-characterized systems, like the language network or the face recognition system, for which certain localizer paradigms have been shown (in fMRI research) to reliably identify the relevant underlying functional neuroanatomy, obtaining the anatomical information from MEG becomes not critical: we already know where these signals are coming from based on dozens or even hundreds of fMRI studies, which are ideally suited for localizing functions. As a result, we argue for leveraging the advantages that MEG has over other recording modalities, the core one being the ability to study the detailed time-course of information processing. Our proposed method provides a means for the selection of language-responsive sensors in which in a second stage different time-resolved processes contributing to sentence comprehension can be studied. Specifically, the fine temporal resolution of MEG offers the potential to study the incremental construction of sentence structure and meaning in real time (e.g., (Heilbron et al., 2022; Ten Oever et al., 2022; Desbordes et al., 2024)). Previous MEG studies have suggested that the temporal dynamics of sentence processing entail both feed-forward as well as recurrent processing (Hultén et al., 2019), that representations extracted from artificial neural network language models capture some aspects of neural signals recorded with MEG (Choi et al., 2021), and that different frequency bands may reflect distinct cognitive processes, such as lexical retrieval, semantic composition, and prediction of upcoming words (Lam et al., 2016; Prystauka & Lewis, 2019).

While the main purpose of this work is not to make inferences about the functional role of each frequency band during language comprehension, we do find that the results align with current research. For example, the observed increase in theta-band power during the sentence condition (Figure 3) supports leading theories proposing that theta synchronization plays a role in retrieving incoming words from memory during sentence processing (Meyer et al., 2015; Meyer, 2018; Prystauka & Lewis, 2019). Similarly, the reported increase in beta-band power aligns with accounts suggesting that beta-band synchronization supports the maintenance of the ongoing cognitive set, facilitating top-down predictions (Lam et al., 2016; Armeni et al., 2019; Prystauka & Lewis, 2019; Lewis et al., 2023). On an additional note, we report differences in the SOIs that are selected at the level of the frequency bands (and the time domain by extension). Sensors that play a key role in one frequency band, might not be equally influential in other frequency bands, highlighting the difference in functional roles that these bands have during language comprehension. This aligns with findings showing that the language network that we can measure with MEG is not driven by a single dominant frequency but rather exhibits a frequency-dependent organization (Coolen et al., 2020).

In summary, the functional identification MEG approach presented here can increase the sensitivity and interpretability of future investigations of the temporal and spectral dynamics of language processing, as needed to decipher the precise computations that enable language comprehension.

### 4.3 Limitations

As is typical with MEG, we used neural changes during sentence processing relative to the baseline to identify the SOIs rather than a contrast between two conditions. In the fMRI implementation of the language localizer, the cortical regions of interest are identified by the sentences > nonwords univariate contrast. It is known that the inverse contrast (nonwords > sentences) activates the multiple demand network (Duncan, 2010; Fedorenko et al., 2013) which is neuroanatomically in close proximity to the language network (Blank et al., 2014; Braga et al., 2020; Du et al., 2024). The inherently lower spatial resolution of MEG may result in sources from the language and multiple demand networks being captured by the same SOIs. Here, we opted to select SOIs based on the sentence condition and verified that the response to the nonwords condition was significantly lower from the sentence condition. The advantage of using the contrast of the sentence minus the nonwords condition is that non-language processing is subtracted out: in the case of the nonwords condition, this processing includes working memory and visual perceptual processing. As we did not use a contrast and examined temporal aggregates within the post-stimulus time window – in which both visual processing and language processing occurs – some of the selected sensors may reflect sources in the occipital cortex related to lower-level visual processes. However, it is clear that most SOIs are not occipital channels (Figure 2).

Secondly, we opted to include the top 10% of most responsive sensors in the sentence condition. This choice assumes an event related synchronization in each of the neural frequency bands over the course of the sentences. While past research does point in this direction (Prystauka & Lewis, 2019), some have demonstrated opposite effects in which, depending on the anatomical brain region, the alpha and beta band also exhibit an event-related desynchronization (Lam et al., 2016).

Thirdly, we opted to compute the percentage signal change from baseline between the fifth (non)word and the end of the trial to calculate the effect size. This choice did not critically impact the results: similar findings were observed when including signals across the whole trial (Supplementary Tables 1 and 2).

Finally, extracting frequency information from the signal can pose challenges. For one, the detectability of neural activations in higher frequency bands (gamma) is limited in MEG (Jerbi et al., 2009; Muthukumaraswamy, 2013). The lower signal-to-noise ratio for high-frequency activity might explain why the within-participant correlations and the effect sizes were smaller in the gamma bands when compared to the lower-frequency bands (theta, alpha, and beta). Additionally, the paradigm used in this study, due to its time-locked stimulus presentation, is expected to give rise to both evoked responses as well as induced responses. A relevant question related to the earlier notion that the frequency analysis presents a different yet complementary perspective to the time-domain analysis, is that the power-based effect sizes in the frequency domain may be driven by the same evoked responses that shape the results in the time domain. To address this, we performed a control analysis in which we estimate the induced power in each of the frequency bands after subtracting the event-related average from the signal. Our verification analysis (Supplementary Figure 1) confirms that the power-based effects primarily reflect modulations of ongoing oscillations.

## 5. Conclusion

Using an extensively validated fMRI language localizer task, based on sentence reading, we showed that the neural responses recorded with MEG are reproducible at the individual participant level in the time and frequency domain and we generalized these findings across two datasets in two different languages (English and Dutch). We observed that language-responsive sensors are spatially variable across individuals, giving an individual-level approach an advantage over the traditional group-level analysis. This method has a wide range of applications—from the detailed characterization of the time-course of language processing to probing the language network in (small) non-neurotypical populations – and may generally encourage the use of MEG to study the functional neuroanatomy of human higher-order cognition.

## 6. Data and Code Availability Statements

Initial preprocessing of the data made use of MaxFilter version 2.2, while further processing was performed with MNE-Python version 1.8.0. All scripts used to generate the outputs for this work as well as the task stimuli are available on OSF (https://osf.io/vc2bw/). Raw MEG data can be made available upon reasonable request.

## Acknowledgements

The authors wish to thank dr. Anna Ivanova and dr. Antonin Rovai. They also acknowledge the Athinoula A. Martinos Imaging Center at the McGovern Institute for Brain Research. This work was supported by an FWO research grant to RB (1509318N), an FWO junior and senior postdoctoral fellowship to RB (12I2117N, 12I2121N), an FWO travel grant for a short stay abroad to RB, and a MISTI Belgium seed fund to EF and RB. RB and MH are supported by the Francqui Foundation. AP was supported by a diversity supplement to EF’s NIH grant R01-DC016950. EF was additionally supported by NIH awards R01-DC016607, R01-DC016950, and U01-NS121471, and by research funds from the McGovern Institute for Brain Research, the Brain and Cognitive Sciences department, and the Simons Center for the Social Brain.

## CRediT contributions

Conceptualization: RB and EF

Methodology: MH, RB, AP, MSi, VW, IB and EF

Software: MH, RB, AP, MSi, CS

Investigation: MH, RB, AP, CS, BL, MSi, VW, MSj, IB, SG, XDT, EF

Data curation: MH, RB, MSi, AP, BL

Formal analysis and validation: MH, RB

Visualization: MH, RB

Writing-original draft: MH, RB, EF

Writing-review and editing: CS, SG, MSj, IB, VW

Supervision and project administration: RB and EF

## Supplementary materials

**Supplementary Figure 1:**
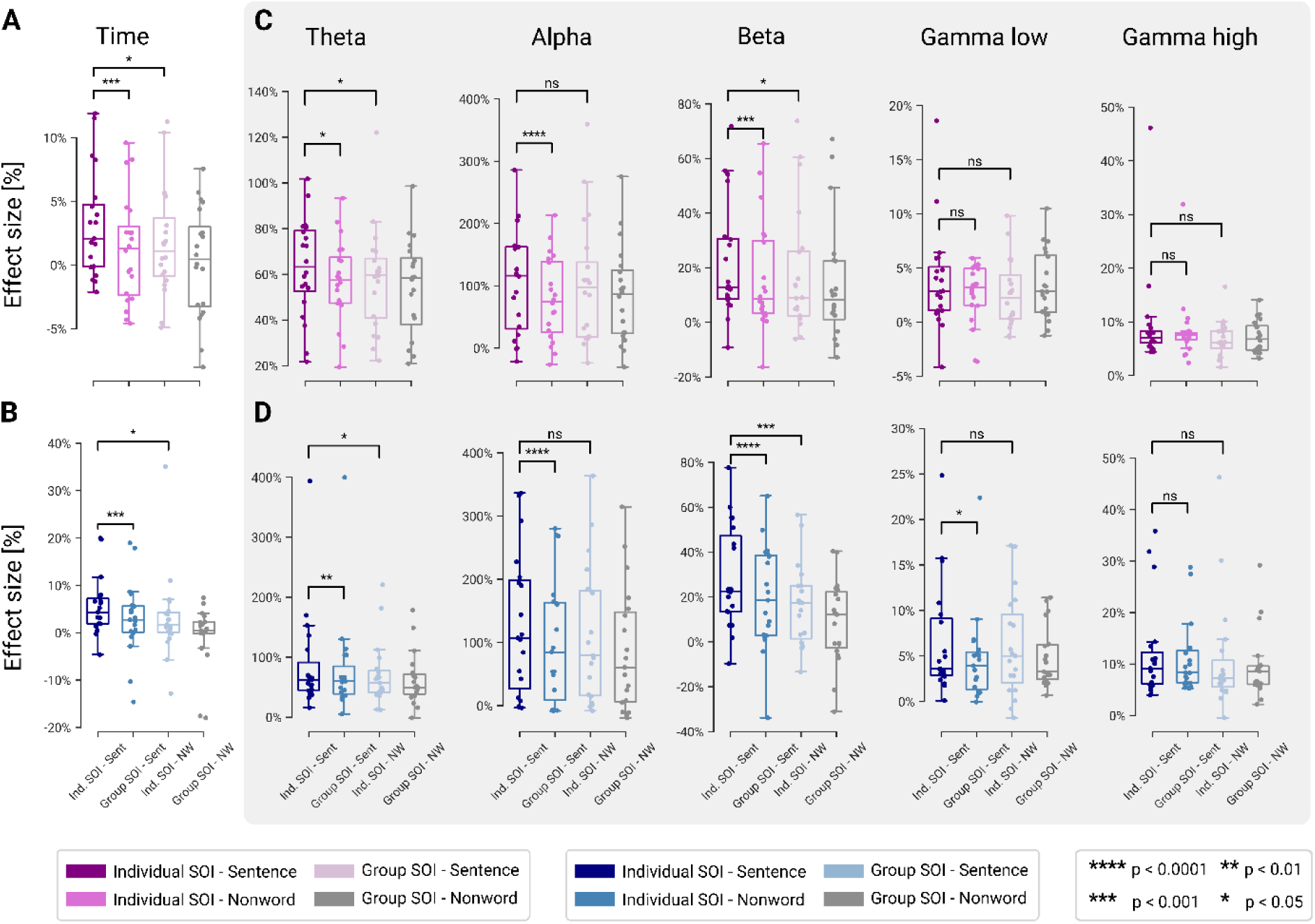
Induced power effect sizes. ***Supplementary Figure 1:*** *Mean effect sizes (percent signal change) for the sentence and nonwords conditions when the language-responsive sensors of interest (SOIs) are defined at the individual level vs. at the group level in the A) English and B) Dutch datasets for the time domain analysis. Similar results depicted for each of the neural frequency bands in the frequency domain analysis after subtraction of the event-related average in the C) English and D) Dutch dataset. In all cases, the SOIs are defined using one half of the data (odd-numbered trials) and the response magnitudes are examined in the other half of the data (even-numbered trials)*.

**Supplementary Table 1:** Within vs between participants correlations in English dataset from start word 1. *Average within participants correlation values and between participants correlation values across odd and even-numbered trials when trials are analyzed from the first word onwards instead of from the fifth word and onwards*.

| **English** | Time | Theta | Alpha | Beta | Gamma low | Gamma high |
| --- | --- | --- | --- | --- | --- | --- |
| Within | 0.58 | 0.59 | 0.86 | 0.77 | 0.55 | 0.13 |
| correlations | (s.d. 0.23) | (s.d. 0.16) | (s.d. 0.10) | (s.d. 0.13) | (s.d. 0.20) | (s.d. 0.13) |
| Between | 0.12 | 0.12 | 0.46 | 0.38 | 0.20 | 0.02 |
| correlations | (s.d. 0.23) | (s.d. 0.21) | (s.d. 0.19) | (s.d. 0.24) | (s.d. 0.20) | (s.d. 0.10) |
| P value | P < 0.0001 | P < 0.0001 | P < 0.0001 | P < 0.0001 | P < 0.0001 | P < 0.001 |

**Supplementary Table 2:** Within vs between participants correlations in Dutch dataset from start word 1. *Average within participants correlation values and between participants correlation values across odd and even-numbered trials when trials are analyzed from the first word onwards instead of from the fifth word and onwards*.

| **Dutch** | Time | Theta | Alpha | Beta | Gamma low | Gamma high |
| --- | --- | --- | --- | --- | --- | --- |
| Within | 0.64 | 0.67 | 0.84 | 0.78 | 0.41 | 0.20 |
| correlations | (s.d. 0.20) | (s.d. 0.15) | (s.d. 0.13) | (s.d. 0.13) | (s.d. 0.21) | (s.d. 0.17) |
| Between | 0.25 | 0.23 | 0.47 | 0.48 | 0.13 | 0.06 |
| correlations | (s.d. 0.25) | (s.d. 0.21) | (s.d. 0.17) | (s.d. 0.19) | (s.d. 0.17) | (s.d. 0.11) |
| P value | P < 0.0001 | P < 0.0001 | P < 0.0001 | P < 0.0001 | P < 0.0001 | P < 0.001 |

